# Bleb formation induced by acidic mixing buffers improves liquid stability of mRNA-LNPs

**DOI:** 10.64898/2026.03.05.709631

**Authors:** Julian Grundler, Beata Chertok, Anoop Nilam, Anna Edmundson, Mason Song, Michael Newton, Matthew R. Scholfield, Adora M. Padilla, Nicole M. Payton

## Abstract

mRNA-lipid nanoparticles (LNP) have proven their potential as a rapidly adaptable vaccine platform and promise to revolutionize numerous therapeutic areas. A major hurdle towards the widespread adoption of mRNA-LNP vaccines and therapeutics is their limited liquid shelf-life compared to more established modalities currently necessitating an ultralow temperature cold-chain to enable their distribution and storage. While ongoing efforts aim to improve liquid stability through chemical modification of mRNA and lipid components, complementary strategies that are broadly applicable across chemistries may further accelerate translation. Here, we present an approach to improve the liquid shelf-life of mRNA-LNPs that does not rely on modifications to the mRNA or LNP chemistry. In particular, we show that bleb formation induced by high ionic strength acidic citrate buffers during LNP formation reduces mRNA degradation and retains in vitro activity during extended liquid storage. We observed an increase in the in vitro activity storage half-life from 2.8 to 18.9 days at 25°C when prepared using high ionic strength buffers translating into a ∼7-fold improvement in the liquid shelf-life of MC3-LNPs. This enhanced stability of LNPs with large amount of bleb formation was mainly attributed to reduced rates of lipid-mRNA adduct formation and mRNA fragmentation. Furthermore, the acidic buffer dependent stabilization was observed across different ionizable lipids with the extent dependent on the ionizable lipid head group. We envision that the induction of bleb formation via selection of appropriate acidic mixing buffers may represent a universal approach to enhance mRNA-LNPs stability and enable extended long-term refrigerated storage.

---

The rapid development of effective mRNA-LNP vaccines during the COVID-19 pandemic demonstrated the advantages of mRNA-LNP drug products compared to conventional biologics by establishing a platform that can be quickly modified to adapt to emerging viral strains.^1^ Specifically, this platform approach reduces product specific development and validation activities resulting in faster to market products while offering adaptability to applications beyond viral vaccines including in vivo expressed biologics, bacterial vaccines, and gene editing.^2–4^ In addition, an established mRNA-LNP platform unlocks the potential for highly personalized therapies by incorporating mRNA sequences tailored to the patient as recently shown in a clinical trial for an mRNA-LNP cancer vaccine that encodes for 20 neoantigens specific to a patient’s tumor.^5^ Despite these benefits, a common shortfall of mRNA-LNP drug products is their limited thermostability. mRNA-LNP vaccines during the COVID-19 pandemic required an ultralow temperature cold-chain with limited liquid shelf-life, restricting their distribution to high-income countries with appropriate storage and shipping infrastructure.^6^ While additional development work has resulted in updated formulations with improved refrigerated shelf-lives, further thermostability improvement is essential to compete with conventional vaccines in a non-pandemic or seasonal setting which can often be stored at 2-8 °C for at least 2 years.^7,8^ Furthermore, the typically shelf-life of standard drug products outside of vaccines typically ranges between 1 and 5 years highlighting the need for novel approaches to improve the stability of mRNA-LNPs to enable competitive mRNA-based therapeutics.^9^

Intraparticle mRNA degradation occurs by both fragmentation and lipid-mRNA adduct formation.^10^ A common strategy to improve the stability of liquid drug products is the optimization of the storage buffer.^11^ Tris buffers at neutral or slightly basic pH have proven to reduce lipid-mRNA adduct formation and enhance freeze-thaw stability, and are the standard in FDA-approved mRNA-LNP products.^7,12^ Beyond storage buffer optimization, modifications to the mRNA structure and sequence can have a major impact on the rate of mRNA degradation.^13^ Stewart-Jones et al. developed a domain-based mRNA vaccine that only encodes part of the COVID-19 spike protein to achieve a thermostable product.^14^ The Kool group demonstrated a method for the reversible acylation of the 2’-OH group that considerably reduced mRNA fragmentation.^15,16^ However, modifying the mRNA sequence or chemistry may compromise mRNA translation and protein expression thus limiting their applicability.^17,18^

LNP-focused approaches mostly rely on changes to the ionizable lipid chemistry. Recently, several groups reported new classes of ionizable lipids to reduce the formation of lipid-mRNA adducts.^19,20^ For example, Hashiba et al. designed a library of ionizable lipids with cyclic piperidine head groups that exhibited significantly improved liquid stability by lowering the amount of reactive aldehyde impurities.^19^ Ripoll et al. circumvented the potential for lipid-mRNA adduct formation caused by the degradation of tertiary amines by using an imidazole head group.^20^ While both of these studies demonstrated enhanced liquid stability, questions remain about the impact of chemical modifications to the ionizable lipid head group on the efficacy and immunogenicity of the resulting mRNA-LNP.

To date, the mRNA-LNP formation process and changes to the LNP morphology have not been thoroughly studied with regards to thermostability despite being potentially applicable across different ionizable lipids and mRNA constructs. Geng et al. showed that optimization of the post-mixing dilution and buffer exchange process resulted in significantly enhanced liquid stability using conventional ionizable lipids with tertiary amine head groups.^21^ The authors further suggested that their improved process may impact the particle fusion mechanism post-mixing leading to differences in LNP morphology as indicated by the reduced amount of empty LNPs.^21^

The importance of the acidic mixing buffer used during mixing of the mRNA with the lipids in ethanol was first demonstrated by the Cullis group who reported that high ionic strength citrate buffers drastically increase the transfection efficiency of resulting LNPs by promoting the formation of mRNA-filled blebs attached to the main LNP core.^22^ Several others have since investigated the effect of acidic mixing buffers on LNP formation and the impact of blebs on transfection efficiency with mixed results.^23–26^ Binici et al. observed that increasing the ionic strength of citrate mixing buffers slightly reduced the in vitro transfection efficiency of SM-102 LNPs without impacting LNP size.^24^ However, Kamanzi et al. showed that individual process steps (i.e., buffer exchange, pH, and ionic strength) affect LNP formation and morphology making it difficult to compare results across different studies.^25^ Despite these efforts, the effect of bleb formation and acidic mixing buffers on mRNA-LNP stability has not been extensively studied. Here, we propose the optimization of acidic mixing buffers as a novel strategy to improve the stability of mRNA-LNPs via the induction of bleb formation.

## Results and Discussion

### High ionic strength acidic mixing buffers retain bioactivity of mRNA-LNPs during liquid storage

We investigated the impact of various acidic mixing buffers with high and low ionic strength containing commonly used buffering species (acetate, citrate, citrate-phosphate) on LNP formation and liquid stability (Figure 1a). We prepared LNPs composed of Dlin-MC3-DMA (MC3) ionizable lipid and standard helper lipids (i.e., DSPC, cholesterol, DMG-PEG_2kDa_) using microfluidic mixing with firefly luciferase (FLuc) mRNA diluted in the respective acidic buffer (i.e., 25 mM acetate pH 4, 300 mM acetate pH 4, 300 mM citrate-phosphate pH 4, 300 mM citrate pH 4, 50 mM citrate pH 3). After LNP formation, the resulting mRNA-LNPs were dialyzed into a 20 mM Tris 10% sucrose pH 7.4 storage buffer and stored at 25°C for up to 4 weeks. LNPs prepared using high ionic strength citrate and citrate-phosphate buffers exhibited an increased hydrodynamic diameter between 100 and 110 nm compared to ∼85 nm for LNPs prepared using low ionic strength mixing buffers (Figure S1) consistent with previous reports about potential particle fusion induced by the high concentration of citrate.^22,25^ LNP sizes remained stable during storage with the exception of 25 mM and 300 mM acetate pH 4 LNPs which showed a gradual increase in hydrodynamic diameter over time.

**Figure 1.**
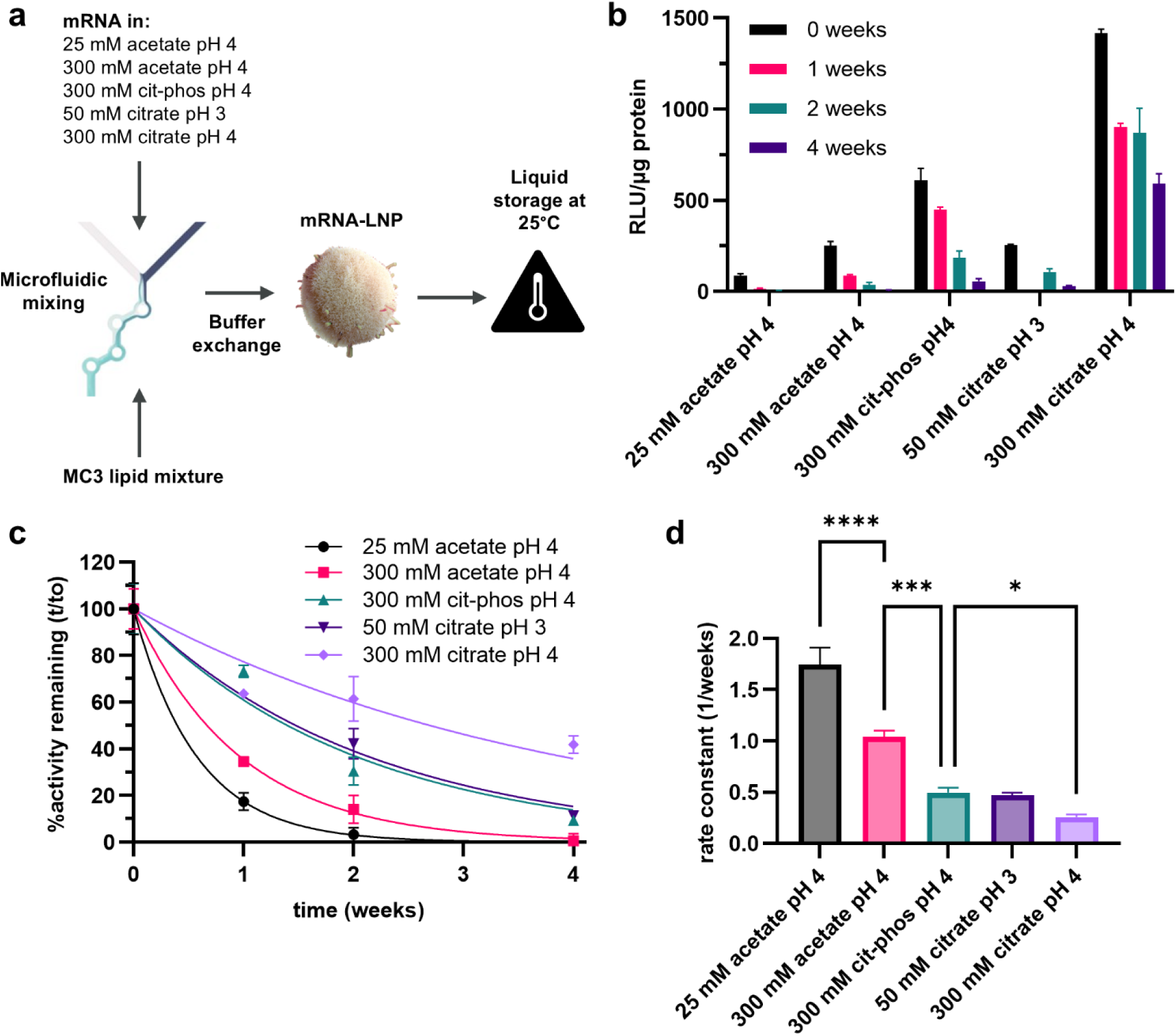
Impact of acidic buffers used during microfluidic mixing of lipids and mRNA on the stability of the resulting LNPs during extended liquid storage. (a) Overview of LNP formation process with lipid mixture dissolved in ethanol containing MC3 ionizable lipid, cholesterol, DSPC, and DMG-PEG_2kDa_, and FLuc mRNA diluted in various acidic buffers. (b) Luminescence of HeLa cells transfected with LNPs that were stored at 25°C for up to 4 weeks. (c) Normalized luminescence of transfected HeLa fitted using a mono-exponential equation. (d) Rate constants obtained from fitting the decrease in LNP transfection potency during 25°C storage. Data are presented as mean ± SD (ANOVA with post-hoc Tukey’s multiple comparison test, *n* = 3). \**P* < 0.05, \*\*\**P* < 0.001, \*\*\*\**P* < 0.0001.

We then monitored the effect of extended liquid storage on the transfection activity of the LNPs by measuring the luminescence of HeLa cells incubated with the LNPs after 0, 1, 2, and 4 weeks storage at 25°C (Figure 1b). High ionic strength buffers increased the initial in vitro activity relative to the 25 mM aetate pH 4 buffer with 300 mM citrate pH 4 outperforming 300 mM citrate-phosphate pH 4 and 50 mM citrate pH 3. The improved in vitro activity of MC3-LNPs prepared using high ionic strength citrate buffer has been previously reported but the impact on stability only received limited attention.^22^ Surprisingly, MC3-LNPs prepared using 300 mM citrate pH 4 buffer not only exhibited high initial activity but also retained ∼50% of their initial activity after 4 weeks of storage at 25°C compared to 25 mM and 300 mM acetate pH 4 buffer LNPs which showed virtually no in vitro activity after only one week of storage (Figure 1b).

This trend was even more evident after fitting the decrease in activity during storage using a mono-exponential equation (Figure 1c). Rate constants obtained from mono-exponential fitting reveal highly significant differences in the in vitro activity decay with increasing ionic strength (25 mM acetate compared to 300 mM acetate) and citrate buffers resulting in smaller decay rate constants (Figure 1d). Notably, the storage in vitro activity half-life increased ∼7-fold from 2.8 days for 25 mM acetate pH 4 MC3-LNPs to 18.9 days for 300 mM citrate pH 4 MC3-LNPs (Figure S2).

### Bleb formation reduces mRNA degradation during liquid storage

We further investigated the mechanism of activity decay during liquid storage. We extracted the mRNA from the LNPs and analyzed the purity using a previously published reverse-phase ion-pairing (RP-IP) HPLC method.^10^ The mRNA purity of MC3-LNPs prepared using 25 mM and 300 mM acetate pH 4 rapidly decreased to ∼20% after one week of storage and no intact mRNA was detectable after two weeks (Figure 2a). In contrast, 300 mM citrate-phosphate pH 4 and 300 mM citrate pH 4 MC3-LNPs still contained ∼50% intact mRNA after 4 weeks closely following the decay in in vitro activity. Furthermore, different mRNA degradation pathways were observed linked to the acidic mixing buffer. The decrease in mRNA purity for MC3-LNPs prepared using acetate buffers can mainly be attributed to the formation of lipid-mRNA adduct during storage (>85% after one week), whereas lipid-mRNA adduct formation is dramatically reduced in 300 mM citrate-phosphate and 300 mM citrate pH 4 buffer MC3-LNPs (Figure S3a) while also exhibiting slightly reduced mRNA fragmentation (Figure S3b). This remarkable difference in mRNA degradation kinetics and mechanism induced by exposure to different buffer conditions is highly unexpected as the acidic buffer is removed during buffer exchange into the neutral pH Tris storage buffer suggesting a structural aspect to mRNA degradation. In particular, high ionic strength acidic buffers have been implicated to affect the LNP morphology and mRNA encapsulation via the formation of bleb structures.^22^

**Figure 2.**
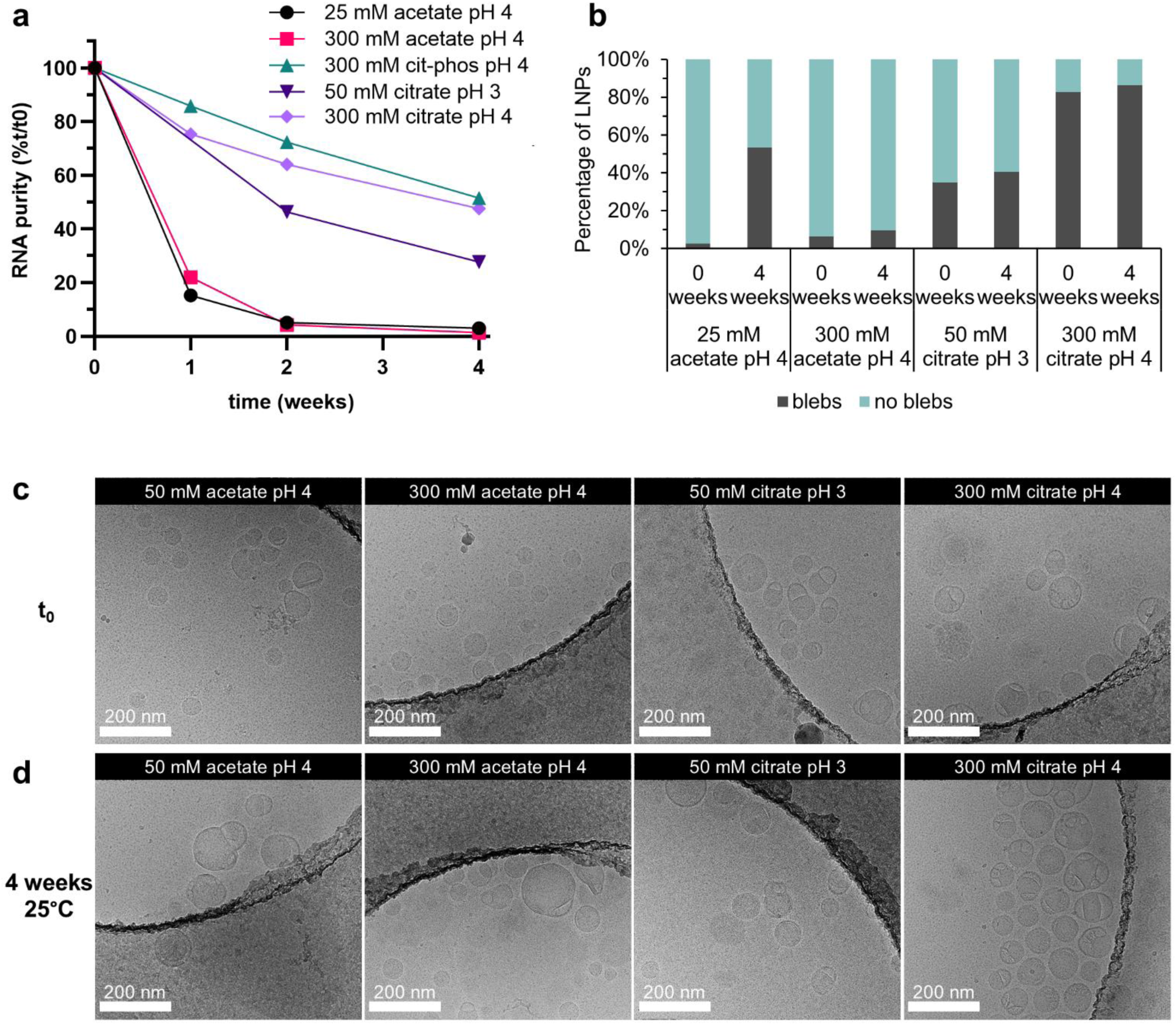
Characterization of MC3 FLuc LNPs prepared using various acidic buffers and stored at 25°C for up to 4 weeks. (a) Normalized purity of mRNA extracted from LNPs assessed via HPLC. (b) Percentage of bleb formation of LNPs before and after storage for 4 weeks at 25°C obtained from cryo-TEM images. Representative cryo-TEM images of LNPs prepared using acetate and citrate buffers (c) before and (d) after storage. Scale bars represent 200 nm.

We tested this hypothesis by assessing the LNP morphology of low and high ionic strength acetate and citrate buffer MC3-LNPs using cryo-TEM. Almost no bleb formation was observed for 25 mM acetate and 300 mM acetate pH 4 buffers before storage (t_0_) with a slight increase at higher ionic strength (Figure 2b and c). However, bleb formation at t_0_ increased to 35% of LNPs containing blebs for 50 mM citrate pH 3 MC3-LNPs and 83% for 300 mM citrate pH 4. Storage at 25°C for 4 weeks resulted in a small increase in blebs for MC3-LNPs prepared using citrate buffers and 300 mM acetate but a much larger increase for 25 mM acetate highlighting that bleb formation can occur at different steps such as during the storage of formed LNPs.^27^ Notably, 300 mM citrate pH 4 MC3-LNPs maintained a high percentage of bleb formation but appear to have more multi-blebbed particles after storage whereas mostly single-blebbed particles are observed before storage (Figure 2c and d).

### Proposed mechanism of mRNA stabilization by bleb formation

Cheng et al. previously speculated that bleb formation is beneficial for intracellular activity by segregating the ionizable lipid from the mRNA thereby stabilizing the mRNA during intracellular processing.^22^ Regular LNPs without bleb formation have been shown to contain the mRNA in aqueous cylinders with the ionizable lipid in close proximity (Figure 3a).^28^ Thus, it is conceivable that the increased interaction between ionizable lipid and mRNA can lead to accelerated mRNA degradation via lipid-mRNA adduct formation and enhanced mRNA fragmentation catalyzed by the amine head group of the ionizable lipid as observed for MC3-LNPs prepared using low ionic strength acetate buffers.^10,29^ In contrast, blebbed LNPs are composed of a hydrophobic “solid core” structure containing the ionizable lipid with the mRNA located inside the aqueous “liposomal-like” bleb compartment. This segregation of the mRNA and ionizable lipid to different compartments effectively minimizes unfavorable interaction during storage resulting in considerably improved stability. Specifically, highly blebbed MC3-LNPs in our study showed drastically reduced lipid-mRNA adduct formation and negligible fragmentation compared to regular non-blebbed MC3-LNPs (Figure S3a and b).

**Figure 3.**
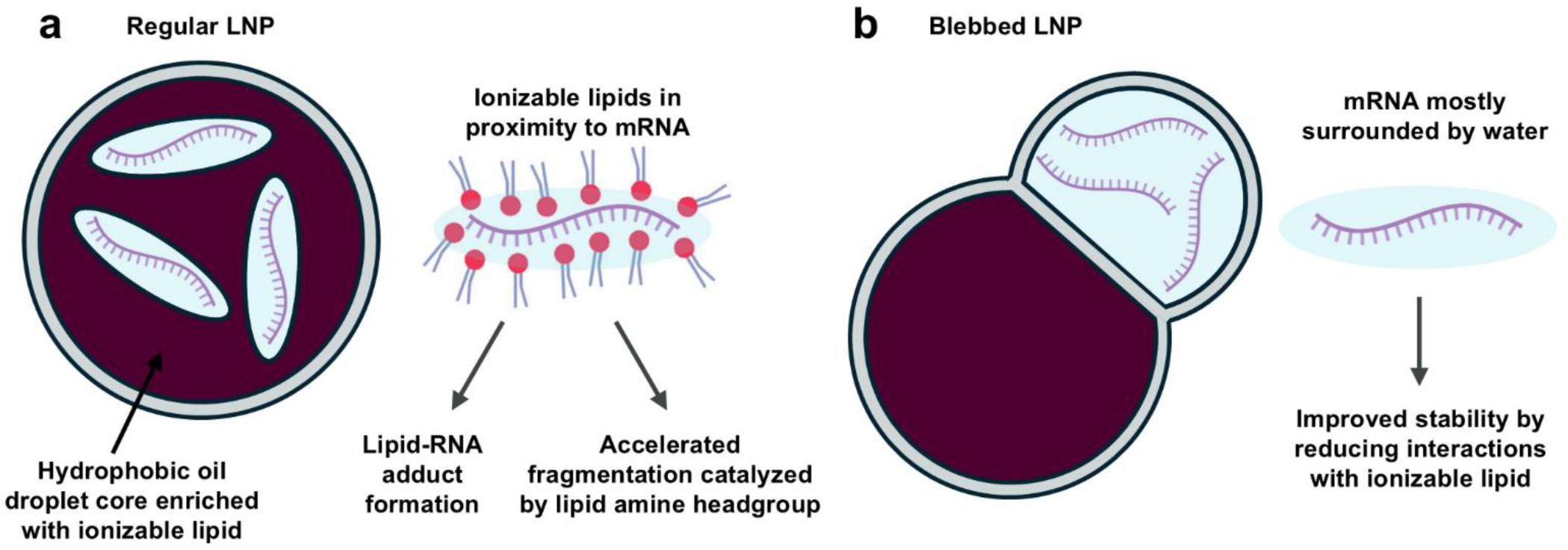
Schematic illustration of proposed mechanism of mRNA degradation inside LNPs. (a) Regular non-blebbed LNP contains mRNA in oil droplet core with ionizable lipids in close proximity resulting in accelerated fragmentation and lipid-mRNA adduct formation. (b) mRNA in blebbed LNPs is mostly contained in aqueous bleb compartment with hydrophobic ionizable lipid located in oil droplet core reducing unfavorable interactions with mRNA during storage.

### Acidic buffer-induced stabilization of mRNA-LNP is affected by ionizable lipid chemistry

The impact of the ionizable lipid on LNP stability and mRNA degradation has been highlighted by several groups.^7,19,30^ In particular, optimization of the ionizable lipid head group was found to lead to major improvements in preserving the activity of LNPs during storage.^19,20^ Thus, we investigated the effect of high ionic strength citrate buffers on the stability of additional LNPs containing various commonly used ionizable lipids (Figure 4a). LNPs were prepared using pH 4 buffers containing either 50 mM acetate, 300 mM acetate, 50 mM citrate or 300 mM citrate and stored at 25°C for 5 weeks. Analysis of the in vitro activity in HeLa cells of LNPs before and after storage reveals that the activity decay is highly ionizable lipid dependent (Figure 4b). Lipid 365 and Lipid 370 LNPs—both containing a 3-hydroxymethyl azetidine head group—retain less than 10% of their initial in vitro activity regardless of the acidic buffers used. Notably, Lipid 370 LNPs prepared using 300 mM citrate buffer exhibited high mRNA purity (∼60% of initial purity) even after 5 weeks of storage at 25°C whereas other buffers led to a purity of less than 30%.

**Figure 4.**
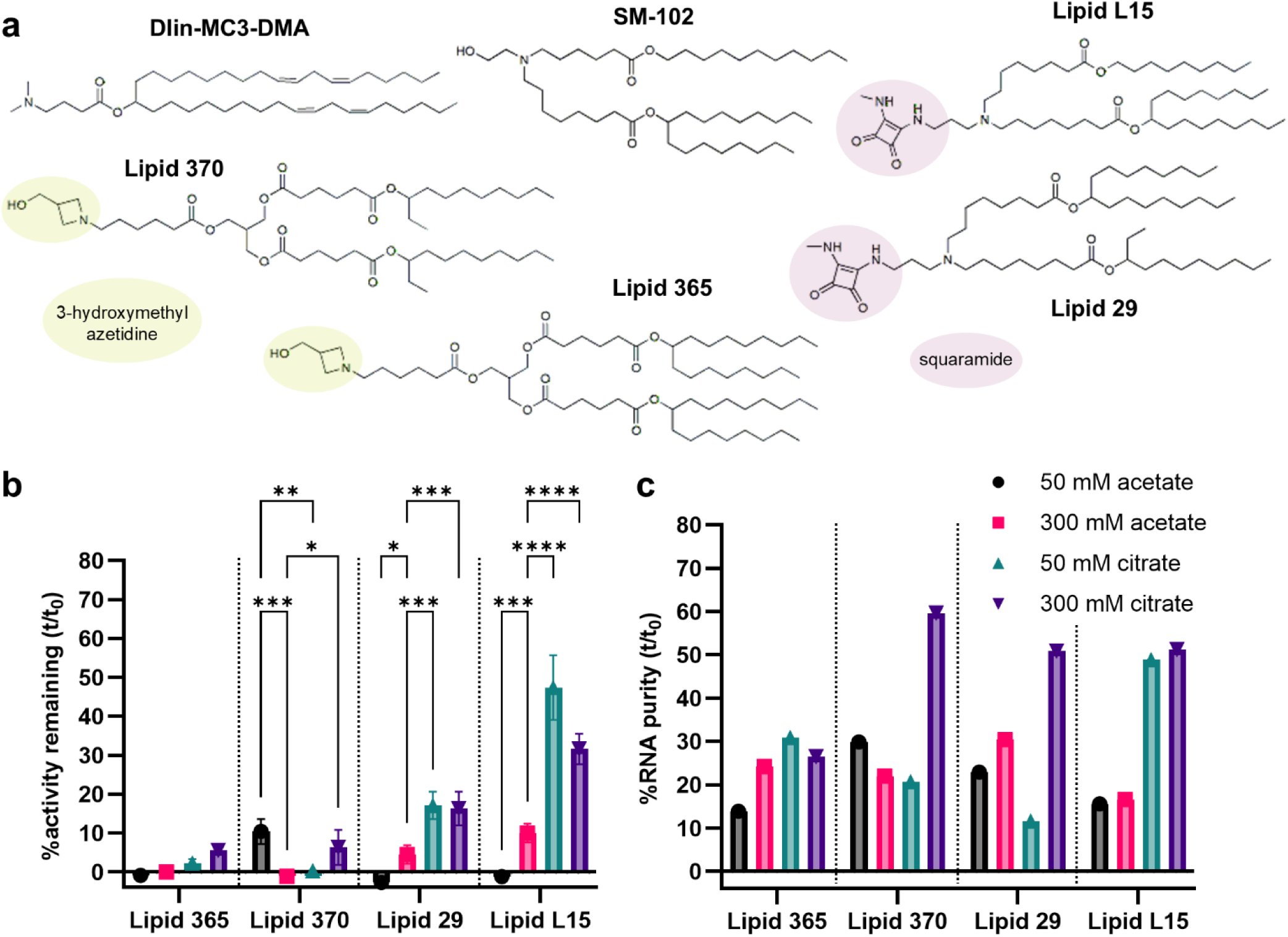
Impact of different ionic strength acidic buffers used to form FLuc LNPs containing various ionizable lipid on their liquid stability during storage at 25°C for 5 weeks. (a) Chemical structures of ionizable lipids used in this study. (b) Ratio of luminescence of HeLa cells transfected with LNPs before and after storage (%activity remaining). (c) Ratio of mRNA purity before and after storage assessed via HPLC. Data are presented as mean ± SD (ANOVA with post-hoc Tukey’s multiple comparison test, *n* = 3). \**P* < 0.05, \*\**P* < 0.01, \*\*\**P* < 0.001, \*\*\*\**P* < 0.0001.

LNPs containing ionizable lipids with squaramide head group (Lipid L15 and Lipid 29) prepared using 50 mM and 300 mM acetate buffers showed negligible in vitro activity (<10%) after storage but exhibited a significant improvement when prepared using citrate buffers (∼20% for Lipid 29 LNPs and as high as 40% for Lipid L15 LNPs). Similar to MC3-LNPs, preparation of Lipid 29 and Lipid L15 LNPs using a 300 mM citrate buffer also resulted in a much higher mRNA purity of ∼50% (Figure 4c) indicating that the stabilization of LNPs induced by high ionic strength citrate buffers is affected by the type of ionizable lipid. In fact, some ionizable lipids have been reported to results in bleb formation even in low ionic strength buffer as their structure promotes particle fusion.^22^ Furthermore, the discrepancy between the high mRNA purity and low remaining activity for some LNPs (i.e., Lipid 370 and Lipid 29) suggests additional factors that affect LNP stability during storage that are also not reflected in other quality attributes such as size and encapsulation efficiency (Figure S5).

### mRNA-LNP stabilization and bleb formation is highly dependent on buffer species

In addition to evaluating different ionizable lipids, we also assessed the impact of different buffer species on bleb formation and mRNA stability. Specifically, we investigated the effect of less common acidic buffers on mRNA-LNP stability using a LNP containing a clinically relevant ionizable lipid (SM-102) that is used by Moderna as a part of their respiratory vaccine platform LNP.^7^ Here, we prepared SM-102 LNPs using 50 mM pH 4 buffers containing gluconate, tartrate, succinate, malate, citrate, and acetate. SM-102 LNPs measured about 70 nm when prepared in acetate or gluconate buffer, around 80 nm in tartrate or succinate buffer, and up to 90 nm when prepared in citrate or malate buffers; no difference in encapsulation efficiency was observed across buffers (Figure S6a and b). Cryo-TEM analysis of SM-102 LNPs after preparation showed considerable differences in bleb formation dependent on buffering species (Figure 5a and c). Acetate buffer resulted in negligible bleb formation (<20% of LNPs) while LNPs prepared using citrate buffer exhibited close to 100% bleb formation consistent with the trends observed for LNPs based on other ionizable lipids studied herein. Tartrate and malate buffers also led to high bleb formation (between 60 to 70%), whereas gluconate and succinate buffers only induced a moderate number of blebs with 49% and 30%, respectively.

**Figure 5.**
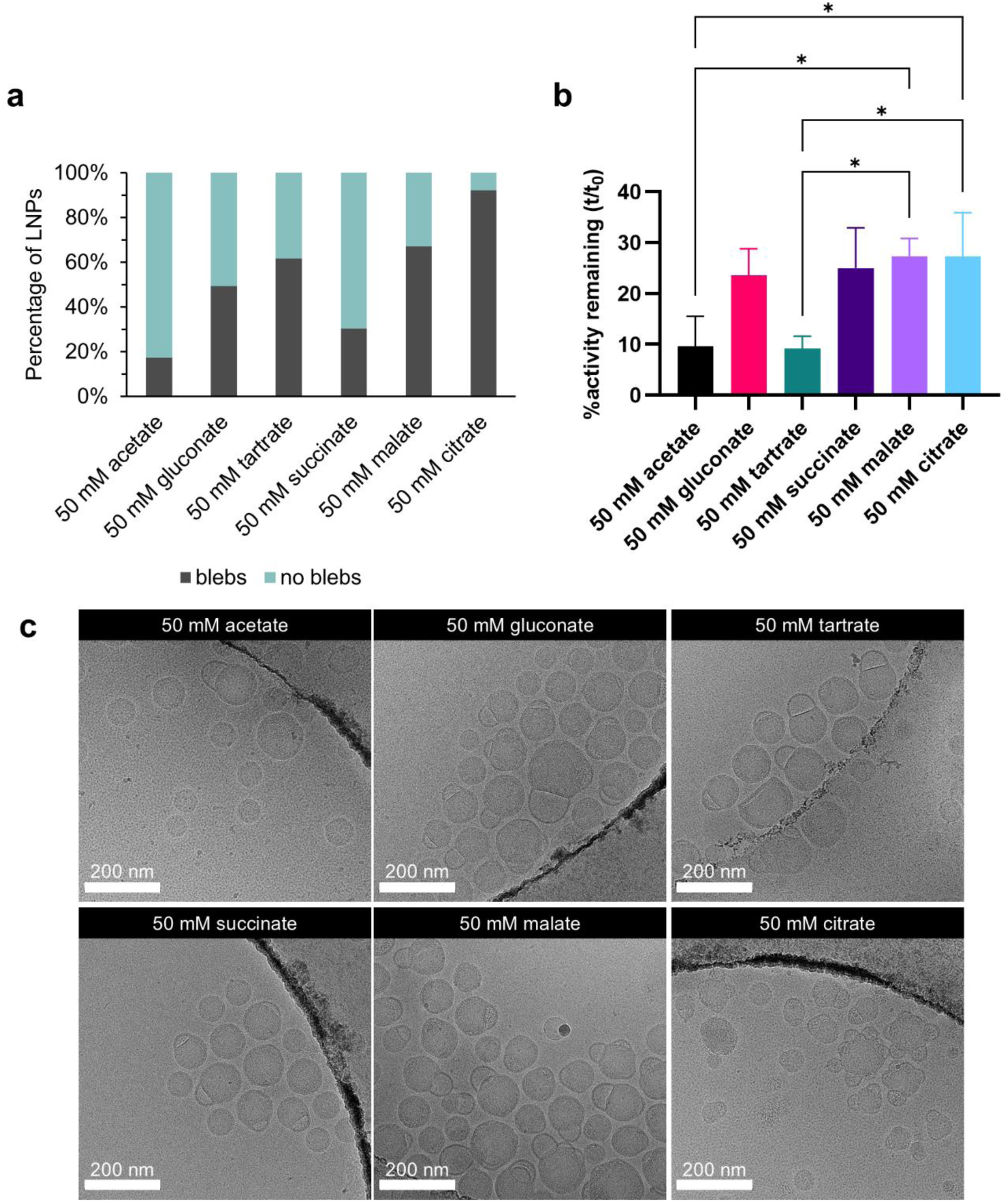
Effect of buffer species of 50 mM acidic buffer used to form SM-102 FLuc LNPs on liquid stability and bleb formation. (a) Percentage of bleb formation of LNPs obtained from cryo-TEM images. (b) Ratio of luminescence of HeLa cells transfected with LNPs before and after storage for 3 weeks at 25°C (%activity remaining). (c) Representative cryo-TEM images of LNPs before storage at 25°C. Scale bars represent 200 nm. Data are presented as mean ± SD (ANOVA with post-hoc Tukey’s multiple comparison test, *n* = 3). \**P* < 0.05.

While it has been previously suggested that citrate buffers increase the fusogenicity of LNPs compared to acetate buffers, limited consideration was given to the differences in charge and apparent ionic strength at similar pH and concentration.^22^ Thus, we calculated the apparent ionic strength of 50 mM acidic buffers at pH 4 and its effect on bleb formation (Figure 6). Generally, a lower apparent ionic strength resulted in reduced bleb formation. However, tartrate—the buffer with the highest apparent ionic strength—led to slightly less bleb formation than citrate suggesting a structural component to fusogenic characteristics beyond apparent ionic strength.

**Figure 6.**
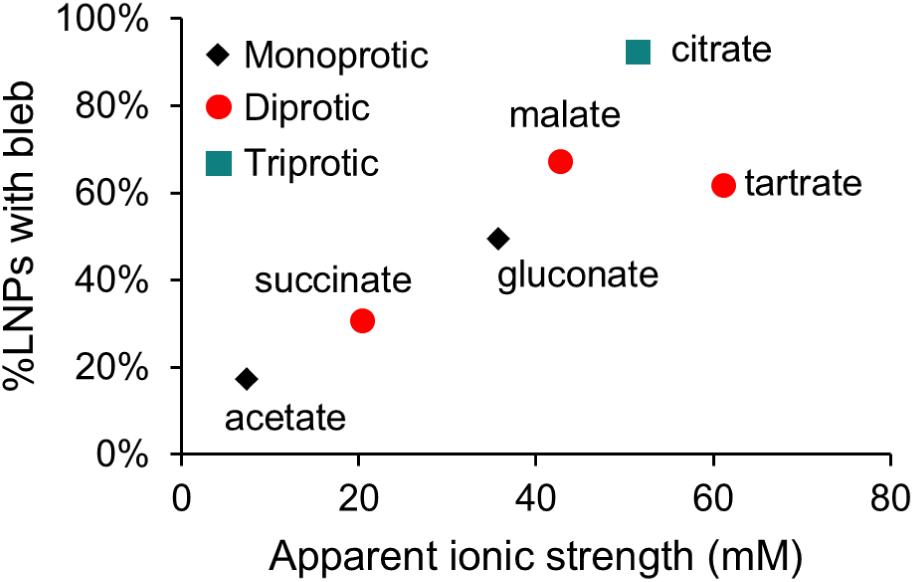
Effect of apparent ionic strength of 50 mM pH 4 acidic mixing buffers on bleb formation of SM-102 LNPs.

Specifically, citrate being the only triprotic acid investigated in this study may facilitate particle fusion due to its unique interaction with the ionizable lipid head group resulting in a bulk phase transition as previously proposed.^31^ Furthermore, Carucci et al. theorized that the buffer specificity can be partially explained using the Hofmeister series.^31^ In our case, citrate, tartrate, and malate being the most kosmotropic ions resulted in increased bleb formation, whereas a limited amount of blebs was observed for more chaotropic ions such as acetate and succinate.^32^

Storage of the SM-102 LNPs at 25°C for 3 weeks resulted in a small increase in size (∼10 nm) irrespective of the acidic buffer used (Figure S6a). Measuring the remaining in vitro activity after liquid storage revealed distinct trends depending on the acidic buffer used to prepare the SM102-LNPs (Figure 5b). Citrate buffer resulted in the highest remaining activity of 27.4% with succinate, malate and gluconate showing comparable results. Acetate and tartrate SM102-LNPs exhibited significantly lower remaining activity compared to other buffers with 9.6% and 9.2%, respectively. This decrease in activity caused by acetate and tartrate buffers coincided with a markedly lower mRNA purity (Figure S7a). In particular, acetate SM102-LNPs only contained 22% intact mRNA compared to 49% for malate SM102-LNPs with the purity loss attributed to both increased fragmentation and lipid-mRNA fragmentation (Figure S7b and c).

While the low activity and mRNA purity observed with acetate was consistent with the limited stability of other LNPs prepared using acetate buffers herein, tartrate—which resulted in high bleb formation—also led to reduced activity and mRNA purity after storage indicating a buffer species specific factor to LNP stabilization beyond bleb formation. It has been previously suggested that buffer species can directly interact and stabilize mRNA inside the LNP.^31^ However, there is conflicting literature about whether buffer molecules from the acidic mixing buffer remain inside the LNP after buffer exchange into the storage buffer or if they will diffuse out of the LNP.^31,33,34^ Thus, additional work is necessary to shed light on the effect of buffer species chemistry on mRNA-LNP stability and potentially identify additional promising buffer systems.

## Conclusion

We demonstrated that mRNA-LNP liquid stability can be influenced by the composition and type of the acidic mixing buffer used during LNP formation. Specifically, we showed that high ionic strength citrate buffers retain the in vitro activity of mRNA-LNPs during extended liquid storage due to the formation of blebs. In addition, the improved stability correlated with a reduced rate of lipid-mRNA adduct formation and mRNA fragmentation suggesting a structural aspect to mRNA degradation. This was further supported by the mRNA degradation being highly dependent on the ionizable lipid head group indicating that the separation of the mRNA inside blebs from the ionizable lipid may prevent unfavorable interactions during storage. Overall, these findings are highly relevant as they elucidate a structural aspect to intraparticle mRNA stability and provide a widely applicable strategy to improve the shelf-life of mRNA-LNPs.

## Methods

Materials and general remarks. FLuc mRNA (5moU, L-7202) was purchased from Trilink, DMG-PEG_2kDa_ from Avanti, DSPC from NOF Corp, and Cholesterol from Sigma-Aldrich. Ionizable lipids SM-102, Lipid 365, Lipid 370, Lipid 29, and Lipid L15 were obtained from Broadpharm and MC3 from Biofine. DLS analysis was conducted by diluting 5 µL of sample in 195 µL PBS followed by analysis using a Zetasizer DLS (Malvern).

mRNA-LNP formation. The ionizable lipid (50 mol%), DSPC (10 mol%), cholesterol (38.5 mol%), and DMG-PEG_2kDa_ (1.5 mol%) were dissolved in ethanol to result in a total lipid concentration of 12.5 mM. FLuc mRNA (1 mg/mL) was diluted in the acidic mixing buffer to a final concentration of 0.12 mg/mL. Mixing of the mRNA with the lipids was performed via a NanoAssemblr Ignite (Cytiva) at an aqueous to ethanol flow rate ratio of 3 to 1 and a total flow rate of 12 mL/min to result in mRNA-LNPs with an N/P ratio of 6. Crude mRNA-LNPs were then buffer exchanged using dialysis cassettes (20 kDa MWCO, Slide-A-Lyzer, ThermoScientific) first against PBS for 2-4 hours followed by dialysis overnight into the final storage buffer (20 mM Tris 10% sucrose pH 7.4). The purified mRNA-LNPs were concentrated using ultracentrifugal filters (30 kDa MWCO, Amicon, Millipore) and a Ribogreen assay was performed to determine concentration and encapsulation efficiency followed by dilution to an encapsulated mRNA concentration of 0.1 mg/mL. All samples were initially frozen and stored at -80°C until the start of the corresponding study.

In vitro cell culture and luciferase assay. HeLa cells (ATCC) were cultured in DMEM medium containing 10% FBS and 1% 10,000 U/mL penicillin-streptomycin at 37°C in 5% CO_2_ humidified atmosphere. The cells were then detached using TrypLE (Gibco), seeded in a 96-well plate (10,000 cells per well), and incubated overnight. The next day, mRNA-LNPs were diluted in DMEM growth medium and 100 ng added per well. After additional incubation for 24 hours, the DMEM growth medium was carefully removed and 100 uL luciferase assay lysis buffer (Bright-Glo, Promega) was added to the cells. After at least 5 minutes incubation at room temperature, 50 uL of the lysed cell suspension was transferred to a separate 96-well plate and an equal volume of luciferase assay substrate (dissolved in PBS) was added followed by immediate quantification of bioluminescence using a plate reader. Bioluminescence measurements were normalized based on total protein concentration by performing a BCA assay (Pierce Micro BCA, ThermoScientific) on the remaining aliquot of the lysed cell suspension.

RP-IP HPLC. The mRNA purity was assessed via a previously published method.^10^ Briefly, 900 µL of 60 mM ammonium acetate in isopropanol was added to 100 µL of 0.1 mg/mL mRNA-LNPs, vortexed, and centrifuged at 14,000 g for 15 min at 4°C. The supernatant was then carefully removed and the pellet redispersed in 1 mL isopropanol followed by vortexing and centrifugation (14,000 g for 15 min at 4°C). After removal of the supernatant, the pellet was vacuum-dried and redispersed in TE buffer for HPLC analysis. The HPLC was equipped with a DNAPac RP column (4 µm, 2.1 × 100 mm, ThermoScientific) and approximately 3 µg of mRNA was injected per sample.

Cryo-TEM analysis. Cryo-TEM sample preparation and analysis was conducted by NanoImaging Services. mRNA-LNPs were vitrified using holey carbon TEM support grids by applying 3 µL of sample to the grid followed by blotting away with filter paper and immediate vitrification in liquid ethane. Sample grids were stored in liquid nitrogen until analysis via a Glacios cryo-TEM microscope (ThermoScientific) equipped with a Falcon 4 direct electron detector operated at 200 kV.

## ASSOCIATED CONTENT

### Supporting Information

The following files are available free of charge.

Additional RP-IP HPLC data including mRNA purity, lipid-mRNA adducts, mRNA fragmentation; LNP characterization via DLS; mRNA encapsulation efficiency data from Ribogreen assay; in vitro activity half-lives for MC3-LNPs. (PDF)

## AUTHOR INFORMATION

### Author Contributions

**Julian Grundler:** Conceptualization, Methodology, Formal Analysis, Investigation, Resources, Data Curation, Writing - Original Draft, Writing - Review & Editing, Visualization, Supervision. **Beata Chertok:** Conceptualization, Methodology, Formal analysis, Resources, Data curation, Writing – Review & Editing, Supervision, Project administration, Funding acquisition. **Anoop Nilam:** Formal analysis, Investigation, Data Curation, Writing – Review & Editing. **Anna Edmundson:** Formal analysis, Investigation, Writing – Review & Editing. **Mason Song:** Formal analysis, Investigation, Writing – Review & Editing. **Michael Newton:** Writing – Review & Editing, Resources. **Matthew Scholfield:** Writing – Review & Editing. **Adora Padilla:** Supervision, Writing – Review & Editing, Funding acquisition. **Nicole Payton:** Supervision, Writing – Review & Editing, Funding acquisition

### Notes

The authors declare the following competing financial interest(s): J.G., A.N., A.E., M.N., M.R.S. A.M.P., and N.M.P. are employed by AstraZeneca. B.C. was employed by AstraZeneca during the development of this work.

## ACKNOWLEDGMENT

The authors acknowledge NanoImaging Services for the collection of cryo-TEM data. Figure 3 was partially created in BioRender. Grundler, J. (2026) https://BioRender.com/mmb6r3n.

