## Supporting Information for "Bleb formation induced by acidic mixing buffers improves liquid stability of mRNA-LNPs"

Biopharmaceutical Development, AstraZeneca, Gaithersburg, USA

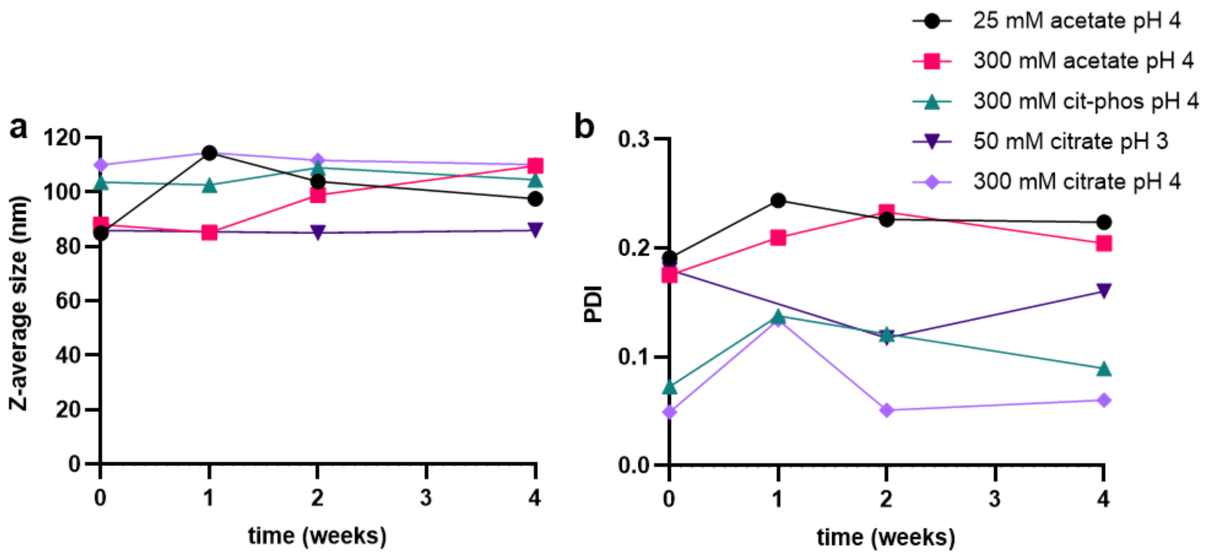


Figure S1. Impact of extended liquid storage at 25°C of MC3-LNPs prepared using various acidic mixing buffer on (a) Z-average size and (b) PDI obtained from DLS.

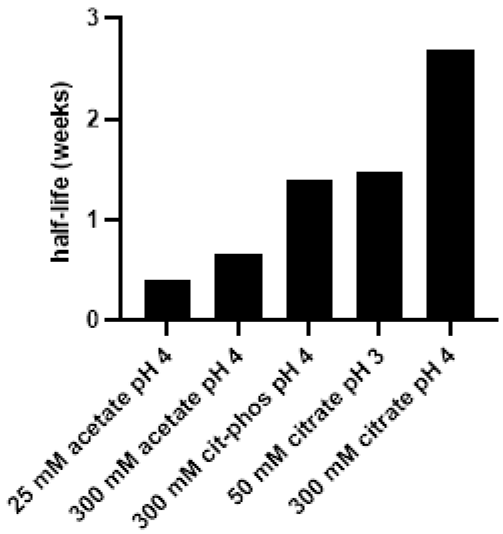


Figure S2. In vitro activity storage half-life of MC3-LNPs prepared using various acidic mixing buffer stored at 25°C obtained from fitting the activity using a first-order decay function.

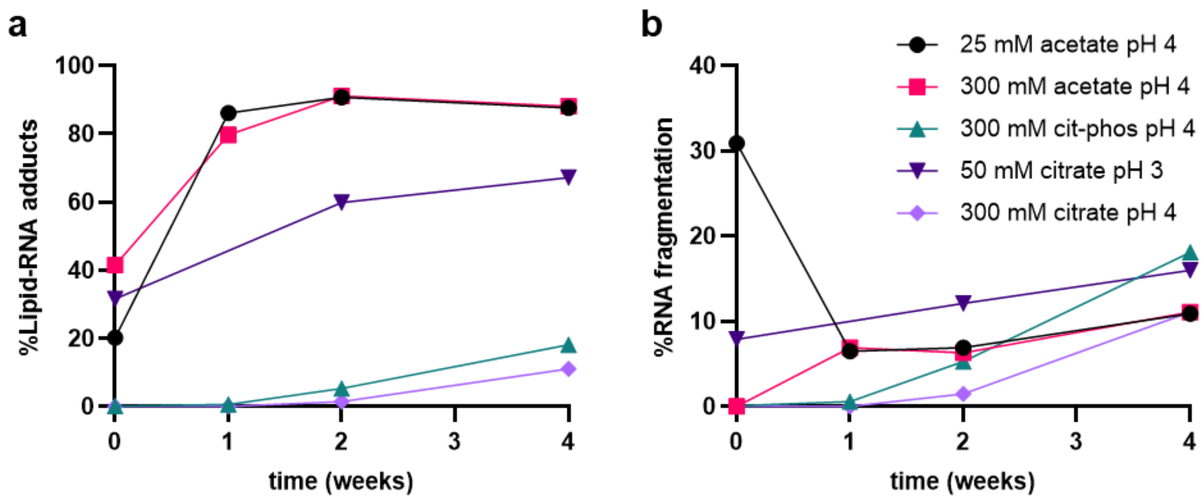


Figure S3. Characterization of lipid-mRNA adduct formation and mRNA fragments of MC3-LNP prepared using various acidic mixing buffers during storage at 25°C obtained via RP-IP HPLC.

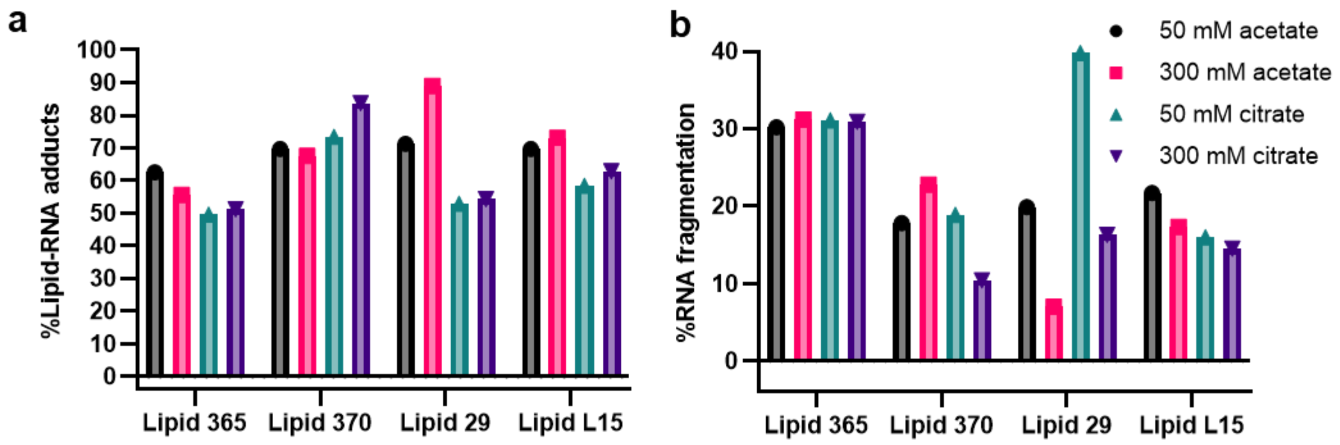


Figure S4. Characterization of (a) lipid-mRNA adduct formation and (b) mRNA fragments of LNPs prepared using various ionizable lipids and acidic mixing buffers during storage at 25°C for 5 weeks obtained via RP-IP HPLC.

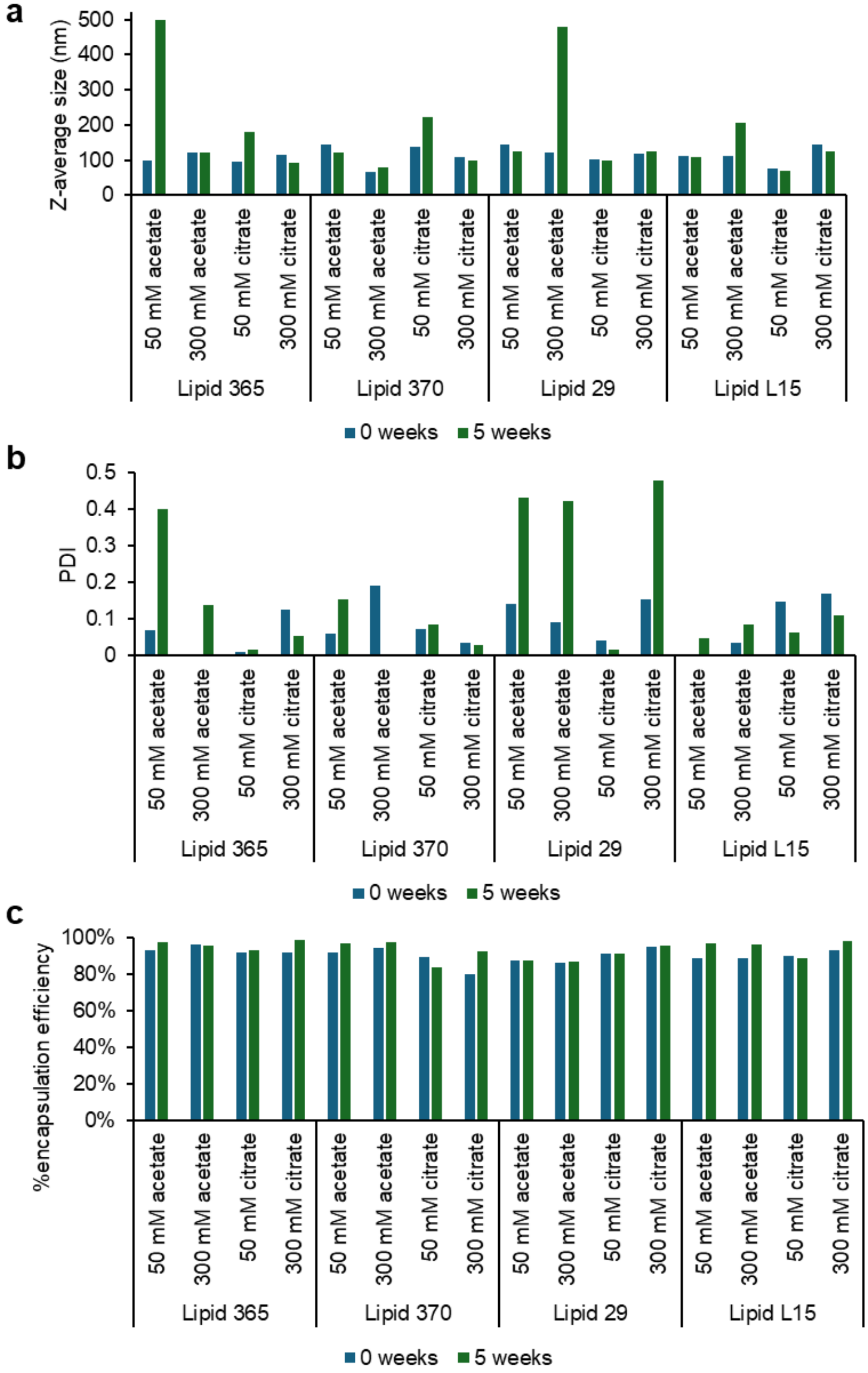


Figure S5. Impact of ionic strength and buffer species of acidic mixing buffer on the attributes of mRNA-LNPs containing various ionizable lipids stored at 25°C for 5 weeks. (a) Z-average size and (PDI) obtained from DLS and (c) mRNA encapsulation efficiency.

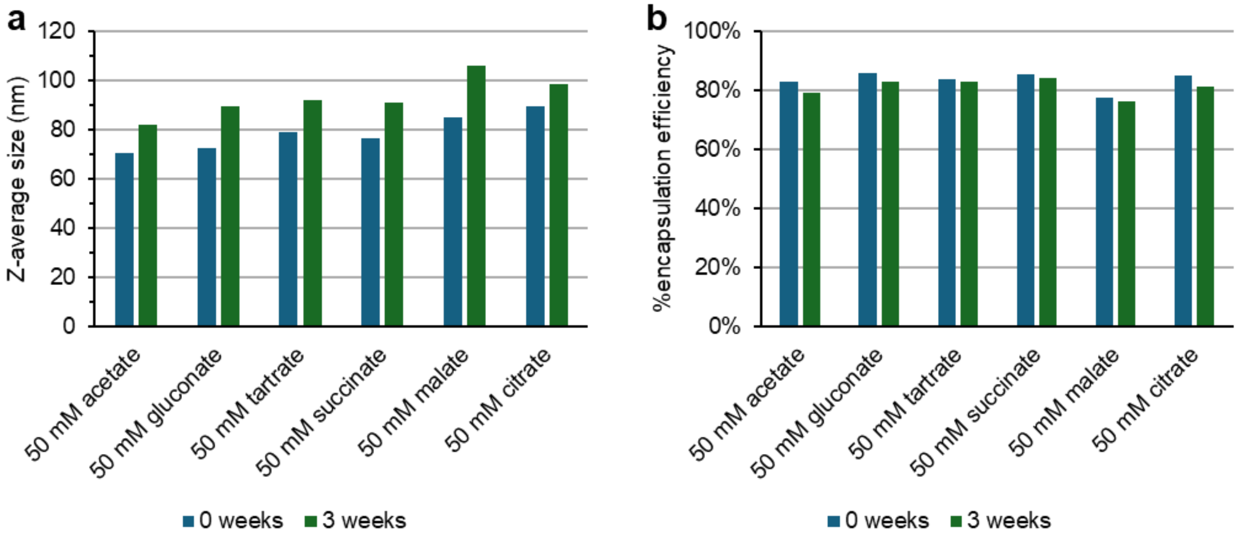


Figure S6. Effect of acidic mixing buffer species on (a) the Z-average size obtained from DLS and (b) mRNA encapsulation efficiency of SM-102 LNPs during storage at 25°C for 3 weeks.

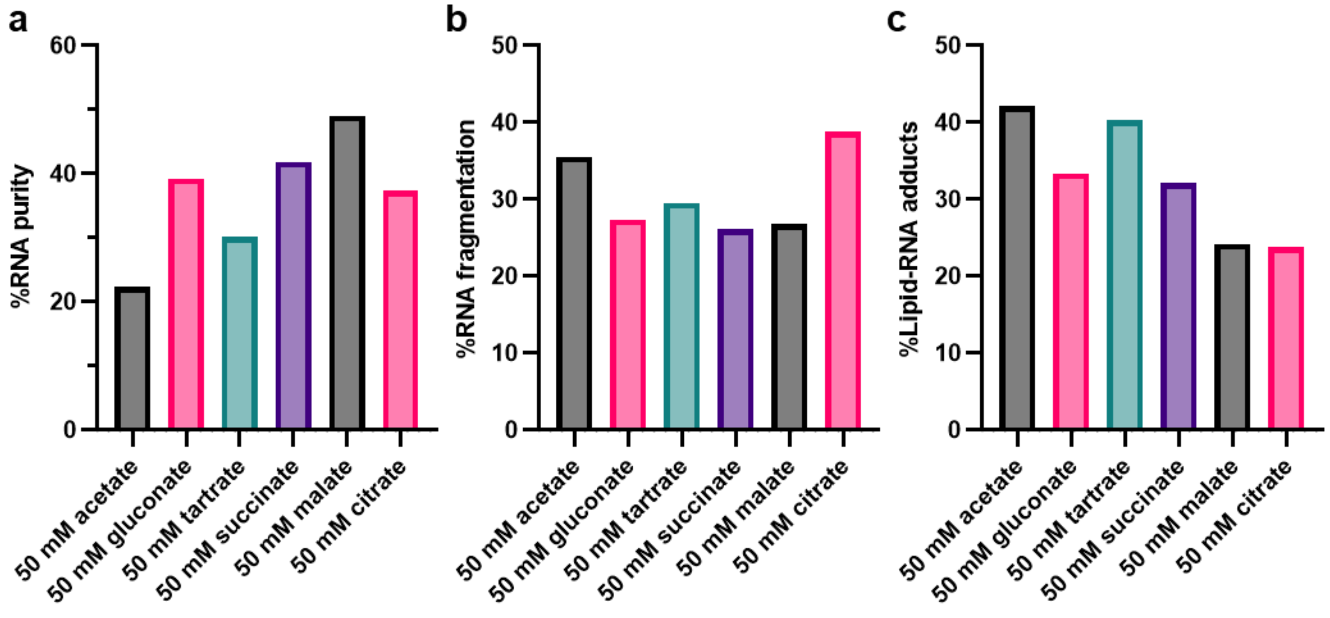


Figure S7. Effect of acidic mixing buffer species on (a) RNA purity, (b) RNA fragmentation, and (c) lipid-RNA adducts of SM-102 LNPs after storage at 25°C for 3 weeks obtained via RP-IP HPLC.

Table S1. mRNA encapsulation efficiency of MC3 LNPs prepared using various acidic mixing buffers.

| sample | Encapsulation Efficiency |
|---|---|
| 25 mM acetate pH 4 | 96.5% |
| 300 mM acetate pH 4 | 97.1% |
| 300 mM cit-phos pH 4 | 96.0% |
| 50 mM citrate pH 3 | 96.8% |
| 300 mM citrate pH4 | 96.4% |
